# Evaluating the utility of identity-by-descent segment numbers for relatedness inference via information theory and classification

**DOI:** 10.1101/2021.09.14.460357

**Authors:** Jesse Smith, Ying Qiao, Amy L. Williams

## Abstract

Despite decades of methods development for classifying relatives in genetic studies, pairwise relatedness methods’ recalls are above 90% only for first through third degree relatives. The top-performing approaches, which leverage identity-by-descent (IBD) segments, often use only kinship coefficients, while others, including ERSA, use the number of segments relatives share. To quantify the potential for using segment numbers in relatedness inference, we leveraged information theory measures to analyze exact (i.e., produced by a simulator) IBD segments from simulated relatives. Over a range of settings, we found that the mutual information between the relatives’ degree of relatedness and a tuple of their kinship coefficient and segment number is on average 4.6% larger than between the degree and the kinship coefficient alone. We further evaluated IBD segment number utility by building a Bayes classifier to predict first through sixth degree relationships using different feature sets. When trained and tested with exact segments, the inclusion of segment numbers improves the recall by between 0.0028 and 0.030 for second through sixth degree relatives. However, the recalls improve by less than 0.018 per degree when using inferred segments, suggesting limitations due to IBD detection accuracy. Lastly, we compared our Bayes classifier that includes segment numbers with ERSA and IBIS and found comparable results, with the Bayes classifier and ERSA slightly outperforming each other across different degrees. Overall, this study shows that IBD segment numbers can improve relatedness inference but that errors from current SNP array-based detection methods yield dampened signals in practice.

## Introduction

Relatedness inference in genetic data often plays a fundamental role in enabling more accurate genetic analyses—both in studies that directly leverage relatives and those that prune them to avoid modeling violations. The need and opportunity to identify genetic relatives continues to increase as the scale of genetic datasets increase^1,2^. One notable example is the UK Biobank wherein roughly 30% of genotyped individuals have a third degree (e.g., first cousin) or closer relative in the study^2^. Applications that make use of genetic relatives are numerous and varied and include pedigree reconstruction^3,4^, pedigree-based linkage analysis for disease and trait mapping^5^, heritability estimation^6,7^, forensic genetics^8^, and genetic genealogy^9^—a popular tool among direct-to-consumer genetic testing customers. On the other hand, traditional genome-wide association study tests and many population genetic models assume that the study samples are unrelated, and, as such, must exclude inferred relatives to avoid spurious signals or inaccurate parameter estimates^10^. All these applications motivate a thorough analysis of the approaches used for relatedness inference to determine which of the various features the methods should leverage.

Many relatedness inference methods only utilize kinship coefficients^11–14^, while some such as ERSA leverage the number of identity-by-descent (IBD) segments between a pair^15^. To date, the question of whether segment numbers provide information for relatedness inference beyond that of kinship coefficients has not been carefully explored. A recent evaluation of 12 pairwise relatedness inference methods using real relatives highlighted three top performing approaches: ERSA and two IBD detection algorithms (i.e., using kinship coefficients derived from their output)^14^. Although ERSA models the distribution of both the number and lengths of IBD segments, that evaluation found that it does not always outperform other methods that only use kinship coefficients. One possible reason is that estimated segment numbers from most phase-based IBD detection methods are inflated due to switch errors that typically break up segments^12,13,16^. Alternatively, these results may indicate that IBD segment numbers and lengths do not better capture relatives’ degrees of relatedness than kinship coefficients.

To determine whether incorporating the number of IBD segments in a model with kinship coefficients (or coefficients of relatedness) improves relatedness inference, we first performed an information theory-based analysis. Feature selection based on information theory is widely used in machine learning and data mining in fields as diverse as bioinformatics and pattern recognition^17–19^. We applied a commonly used measure— mutual information (MI)—to quantify the dependency between various features and the class variable (here the degree of relatedness) and also the dependency among the features themselves. An advantage of this approach is that MI does not make an assumption of linearity between the features and can be calculated for both discrete and continuous variables^20^. In addition, the MI analysis results do not depend on the specific classifier used downstream and can capture the relationship between variables from an information theory perspective that is distinct from classification.

We also conducted a classification-based analysis to determine the importance of IBD proportions and segment numbers for inferring degrees of relatedness. For this purpose, we developed a Bayes classifier with mathematical underpinnings that parallel those of MI. Bayes classifiers are a form of generative learning that seek to minimize the probability of misclassification by estimating the probability of a given data point being from each class^21^. In this work, we assign a pair of relatives to the maximum posterior probability degree, in contrast to approaches that map estimated kinship coefficients to degrees of relatedness using *a priori* fixed ranges of kinship^11,12,14^. The latter ignores the effect of population structure on IBD signals—including background IBD segments^8^. These effects are important to model since they vary by population and can meaningfully influence relatedness classification. Furthermore, bias in the detection of IBD segments can shift the distributions of both IBD proportions and segment numbers. Such biases may especially impact classification of more distant relatives as they have smaller ranges of kinship values that correspond to a given degree. In light of these concerns, we estimate the probability of the features given the degree (i.e., the likelihood) using training data simulated using genotypes from the target population. This implicitly accounts for the influence of background IBD segments as well as any errors in IBD segment detection. Researchers with access to data from a given population can also apply this strategy by using the available samples as founders in simulated pedigrees^22^.

Finally, we benchmarked the performance of our relatedness classifier together with ERSA and IBIS using simulated genotypes. Overall, we obtained comparable classification results for all the methods, indicating that the Bayes classifier is reliable and suggesting that our approach can be used in practice given appropriate training data resources. Notably, the Bayes classifier performs similarly to IBIS (which does not use segment numbers) demonstrating that, in practice, incorporating segment numbers provides very little improvement in classification rates.

All the analyses in this paper leverage IBD segments from simulated data, either exact segments produced by the simulator or segments inferred from simulated genetic data. In particular, we investigated (1) MI quantities based on exact segments, (2) classification rates using exact segments, and (3) classification rates from inferred segments. In this way, the MI analysis quantifies the theoretical information gain available by fully exploiting relatedness signals captured by exact segment numbers. Additionally, the classification analysis using exact segments reveals how much improvement in relatedness inference is possible by incorporating IBD segment numbers in the limit of perfect IBD detection. Finally, comparing the classification results using exact versus inferred segments enables us to localize the influence of IBD detection errors.

## Results

We analyzed the potential for using coefficients of relatedness *r* (defined below) either alone or both *r* and *n*, the number of IBD segments a pair of relatives share, to infer the pair’s degrees of relatedness *D*. To begin, we quantified the inherent dependency between the IBD segment features and *D* by analyzing MI between the features and *D*. MI is a quantification of the information obtained about one random variable through observing another; in this case, we analyzed the information gained about *D* through observing the variables *r, n*, or (*r, n*). We compared our analysis of MI quantities with the corresponding Bayes classification results based on features *r, n*, and (*r, n*); the conclusions we form about the classification effectiveness of different feature sets are therefore based on both the MI and classification results.

Throughout, we refer to IBD regions that two individuals share on only one haplotype copy as IBD1, and those the individuals share IBD on both chromosomes as IBD2.

### Mutual information analysis

We used thousands of relative pairs to estimate mutual information 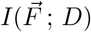 between different IBD features 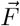 and the degree of relatedness *D* of each pair (Methods, “Estimating mutual information”). Specifically, we compared MI values of *I*(*n* ; *D*), *I*(*r* ; *D*), and *I*((*r, n*) ; *D*) calculated using units of bits. Let 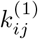 and 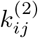 denote the proportion of their genomes that individuals *i* and *j* share IBD1 and IBD2, respectively—i.e., the sums of genetic lengths of all IBD1 or IBD2 segments divided by the total genetic length of the genome analyzed. We calculate *r* as twice the kinship coefficient or 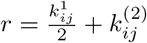.^14^

The first analysis uses exact IBD segments from pairs of individuals that each have one of 13 genetic relationships (Table 1). To reduce the influences of randomness, we replicated this analysis by performing 80 independent simulations. We also analyzed three different distributions of numbers of pairs per degree *D*: uniform, exponential, or a slow-exponential function where the number of pairs increases exponentially with degree for both the exponential and slow-exponential distributions (Figure S1). The exponential function is potentially a more realistic distribution of relatives than the uniform, while the slow-exponential is intermediate between the two.

**Table 1:**
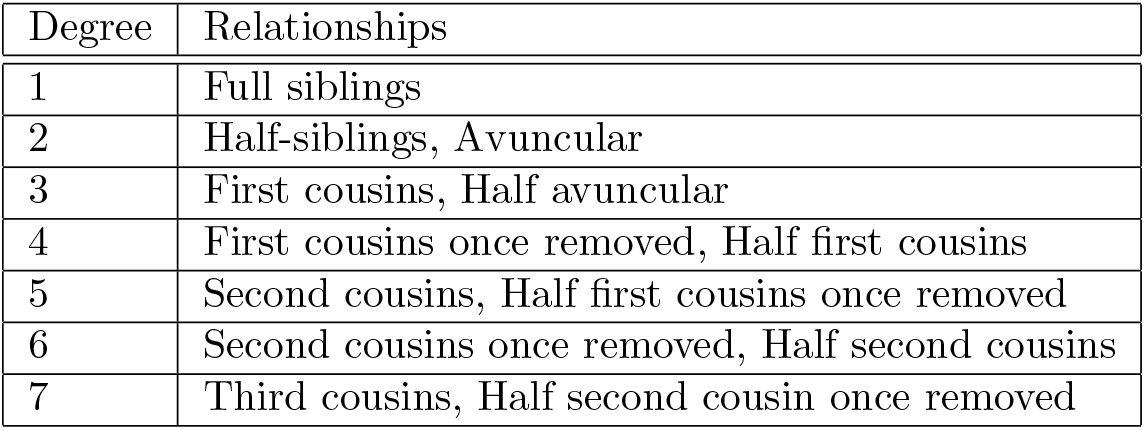
Simulated relationship types in each degree of relatedness. Half relatives share only one common ancestor while other types have two common ancestors.

Figure 1(a) shows the average MI of the simulated pairs computed over all 80 runs (Methods, “Simulated data”). For each distribution shape, the MI between the multivariate feature (*r, n*) and univariate *D* is the greatest, followed by *I*(*r* ; *D*) and *I*(*n* ; *D*). To quantify the relative increase in MI when including both *n* and *r*, we used a normalized MI gain 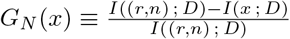 where *x* ∈ *{r, n}*. The normalized MI gain *G*_*N*_ (*r*) (the increase in information gained from using (*r, n*) beyond that of only using using *r*) is 0.030 for the uniform distribution, 0.040 for the slow-exponential, and 0.068 for the exponential. Greater MI indicates a stronger dependency between the features and *D*, and therefore classifying *D* based on features with greater MI should yield greater recall. At *G*_*N*_ (*r*) of 0.068 for the exponential distribution, we expect that incorporating numbers of perfectly detected segments could meaningfully improve classification of degrees of relatedness compared to using *r* alone, especially for higher order degree pairs. In turn, the normalized gain over using segment number alone, *G*_*N*_ (*n*), is 0.15 for the uniform distribution, 0.14 for the slow-exponential, and 0.13 for the exponential, demonstrating that use of *r* dramatically improves classification rates compared to only using *n*, regardless of the distribution of *D*.

**Figure 1:**
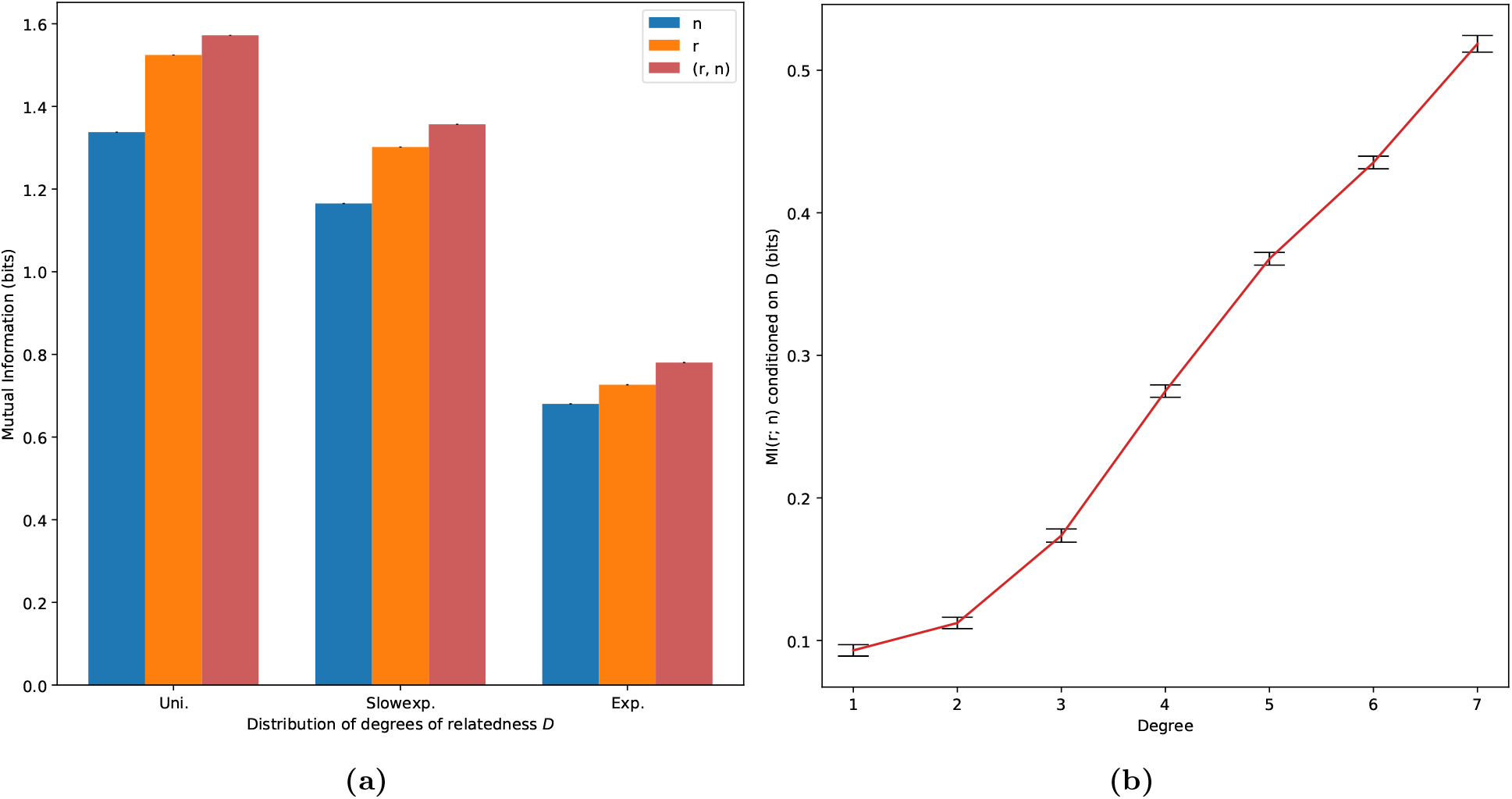
MI between relative pairs calculated using exact IBD segments. MI between (a) various IBD feature sets and *D* and (b) *r* and *n* conditioned on the relatives’ degree of relatedness. All MI quantities are averaged over 80 independent runs, and the values in (b) are calculated using the uniform distribution with 33,000 pairs per degree. Error bars indicate one standard error and are barely visible in (a) (all of order 10^−3^).

Across all three feature sets, the MI is maximal for the uniform *D* distribution and decreases as the distribution becomes more exponential. By construction, the exponential distributions have a higher proportion of high-degree relative pairs compared to the uniform distribution. Therefore, the IBD features from higher degree pairs share less information with *D* than lower degree pairs. This is consistent with observations from classification analyses that show that the recall of degree inference decreases as the degree increases^14^.

To better understand how *r* and *n* relate to each other as well as to *D*, we calculated MI between these two features using the exact IBD segments and conditioned on the degree of relatedness (Figure 1(b)). The amount of shared information between features *r* and *n* monotonically increases with degree of relatedness, meaning that in higher degree pairs *r* and *n* have increased redundancy. Therefore, using both features has less benefit for classification in higher degrees. Nevertheless, both *r* and *n* individually become less informative about *D* with increasing degree, so any additional information can be useful.

### Bayes classification and statistical tests of exact and inferred IBD segments

As MI quantities from exact IBD segments suggest the potential for sizeable improvements by using (*r, n*) to determine *D*, we sought to understand whether parallel results arise from explicit relatedness classification. To that end, we simulated another 210,000 pairs of relatives for training, this time producing genetic data for them using genotypes from UK Biobank unrelated samples as pedigree founders (Methods, “Simulation”). We detected IBD segments in these samples with IBIS and used the resulting *r* and *n* quantities to train Bayes classifiers. For comparison, we further trained a separate set of classifiers using the exact IBD segments from the same simulated pairs (Methods, “Bayesian classification”). Using Bayes classification allowed us to incorporate our prior knowledge of the distribution of *D* to better determine the pairs’ degrees, and also more closely mirrors the mathematical basis of MI. For both the inferred and exact statistics, we generated a set of three classifiers, one trained only on the coefficient of relatedness *r*, one on the IBD segment number *n*, and a third on the vector (*r, n*). We tested both the exact and inferred segment classifiers on 80 independent simulated datasets containing 3,000 simulated relative pairs per degree, again inferring segments with IBIS. (Genetic data for testing pairs was produced identically to the training pairs, as noted above.)

Figure 2 shows the recalls of these classifiers as a function of degree and also shows the recall differences between classifiers trained on (*r, n*) and *r*. We also show the proportions and types of misclassifications in the inferred and exact datasets in Figure S2 and S3. Almost all misclassified pairs are inferred as an adjacent degree of relatedness compared to the truth (i.e., one degree closer or more distant). Note that we do not report accuracy results for seventh degree relatives as these pairs act as an “unrelated” class that provide bounds on sixth degree relatedness classification.

**Figure 2:**
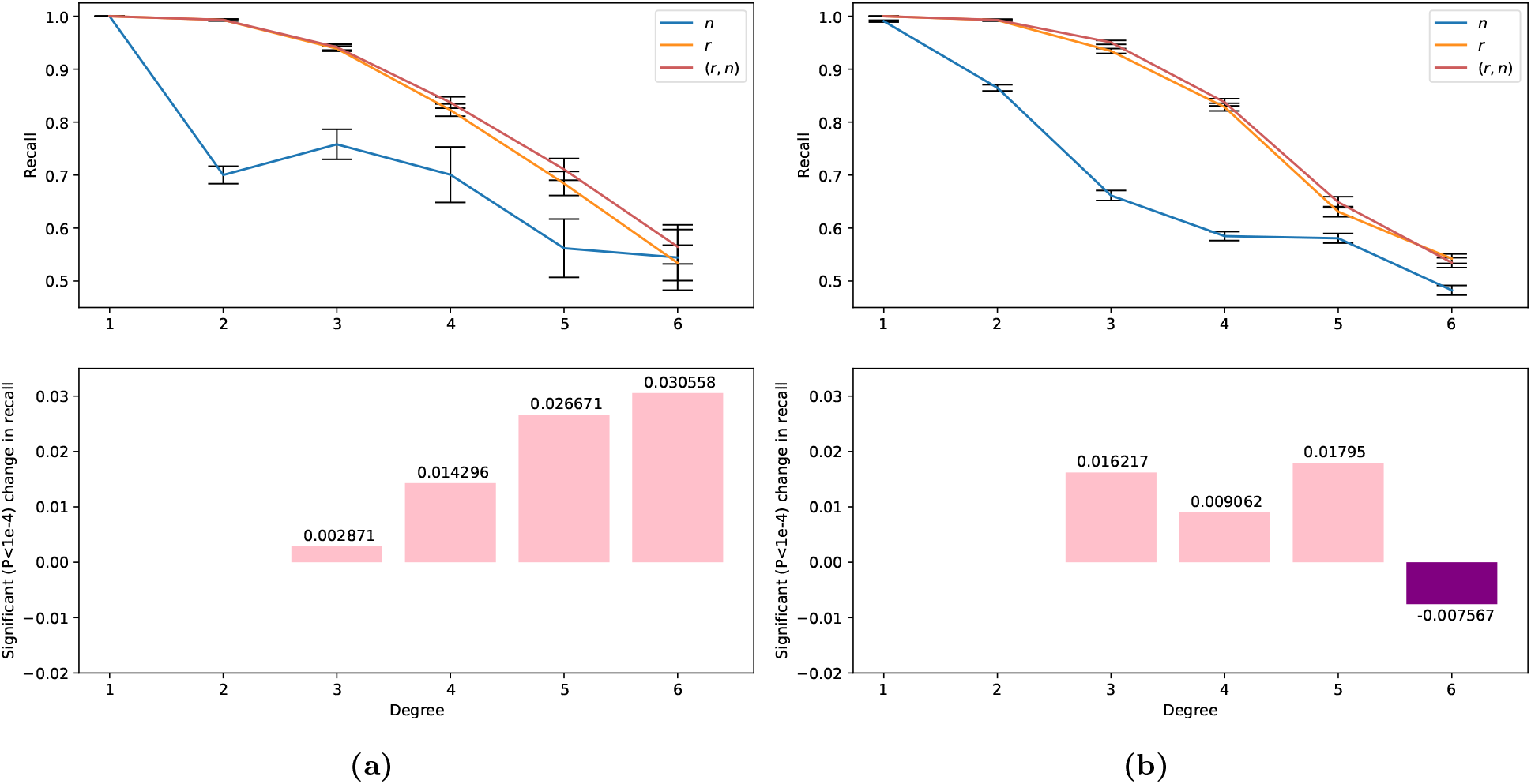
Recalls of Bayes classifiers for first through sixth degree relatives. Results are from classifiers trained on (a) exact and (b) inferred segments with features *n, r*, or (*r, n*). The recalls for both (a) and (b) are calculated using the uniform distribution of 3,000 pairs per degree and averaged over 80 independent runs. For each degree, the lower subplot shows the corresponding significant (*P <* 10^−4^) change in recall between classifiers (*r, n*) and *r* (positive values have greater recall in the (*r, n*) classifier). Significant increases and decreases per degree are shown in pink and purple, respectively. Error bars indicate one standard error.

Overall, recalls for all three classifiers decrease monotonically as a function of the degree of relatedness. For first and second degree pairs, the classifiers trained on *r* and (*r, n*) both have nearly perfect recall values of over 0.99. For higher degree pairs from third through sixth degree, the recalls of the *r* and (*r, n*)-trained classifiers fall from over 0.93 (third degree) to below 0.55. This is consistent with previous observations from real relatives^14^, and aligns well with our results based on MI: The features of higher degree pairs share less information with *D*, meaning that the IBD signals of higher degree pairs tell the classifier less about their true *D* (see misclassification rates in Figure S2 and S3). The classifier trained on *n* alone performs poorly in all but degree one: For second degree relatives, the classifier trained on inferred segments has a recall of only 0.86, and in third through sixth degree relatives its recall is 0.06 to 0.27 units lower than those of the classifier trained on *r*. The results for the classifier trained on exact segments are qualitatively similar to those of the inferred-segment classifier.

In general, when using both exact and inferred IBD segments, the classifiers trained on (*r, n*) outperform those trained on *r* for every degree. One exception is in the inferred IBD segments for sixth degree pairs, where the classifier trained on *r* has a recall of 0.54 while the classifier trained on (*r, n*) has a recall of 0.53. This decrease in recall is counter-intuitive because the (*r, n*) classifier is trained on a strictly larger feature set and so has more information than the *r* classifier. In addition to general stochasticity introduced by segment detection for these distant relatives, it may be that this decrease is caused by the distributions of segment numbers inferred by IBIS (Figures S4 and S5): IBIS does not detect segments smaller than 7 cM and so the distribution of numbers of detected segments for fifth and sixth degree pairs have lower means and are more similar to each other.

We ran two-sided independent sample *t*-tests on the recalls from the (*r, n*) and *r* classifiers trained on the inferred IBD segments. Except for the first degree relatives, in which all three classifiers have recalls of nearly 1.0, and the second degree pairs, in which the two classifiers containing *r* have above 0.99 recall, the differences in recall between the (*r, n*) and *r* classifiers are significant (*P <* 10^−7^) but small in magnitude. These differences range from −0.00756 to 0.0179 in third through sixth degree pairs. In turn, for the classifiers trained on exact IBD segments, the (*r, n*) classifier has significantly greater recall than the *r* classifier in third through sixth degree relatives (*P <* 10^−4^). The improvement in recall ranges from 0.0029 to 0.031, suggesting that better IBD segment inference would meaningfully benefit classification with (*r, n*) (Figure 2).

### Comparison with IBIS and ERSA

To put these results in the context of existing methods, we compared our Bayes classifier with IBIS’s built-in relative classifier and with ERSA, another method that models relatedness using IBD segment number (as well as with segment length). This analysis uses for testing another independent set of 3,000 pairs per degree, again simulated from UK Biobank individuals. Our Bayes classifier remained trained on the same 210,000 pairs as above.

In general, all three methods performed comparably. The accuracy of the Bayes classifier closely tracks that of IBIS, which may be because the Bayes method takes IBIS segments as input. At the two extremes of relatedness we considered, all three methods have similar recalls for first degree and sixth degree relatives with differences smaller than 0.01. The Bayes classifier has nearly identical recall to IBIS in second and third degree pairs (the differences are bounded above by 0.004), whereas ERSA’s recalls for these degrees are 0.06 and 0.02 units smaller, respectively. (An analysis with real relatives also found that ERSA’s second degree classification rates are reduced compared to other approaches^14^.) For fourth degree relatives, ERSA has a recall 0.01 units higher than the Bayes classifier, and 0.035 units higher than IBIS. ERSA also outperformed the Bayes classifier and IBIS on fifth degree pairs by 0.067 and 0.054 units, respectively. ERSA’s improved performance compared to the other two methods may be because of its use of ≥ 2.5 cM segments (instead of ≥ 7 cM segments from IBIS). Consistent with this, simulated fourth and fifth degree relatives have a non-trivial proportion of 3–7 cM segments (Figure S6)—suggesting that these undetected IBD segments may lead to more erroneous calculations by IBIS and the Bayes classifier. Another factor benefiting ERSA is its population model that accounts for background relatedness, which may help it in this and other datasets. Additionally, we used perfectly phased data as input to GERMLINE^23^, and we supplied the resulting segment calls to ERSA (Methods, “Simulated data”). Notably, ERSA’s higher recalls for fourth and fifth degree pairs are close to the range of the Bayes classifier’s recalls using exact IBD segments (in fact, ERSA outperforms the exact Bayes classifier in these degrees by 0.012 and 0.0031, respectively). Finally, considering run time, the Bayes classifier is efficient, taking on average 1 minute 40 seconds to analyze the test data and 7 seconds to train on the 210,000 training pairs. In contrast, ERSA takes more than 3.5 CPU days to classify the testing pairs.

## Methods

### Mutual information discrete definition and binning approaches

MI is difficult to calculate for continuously valued variables without a known distribution and whose distribution must therefore be estimated from finite data. Furthermore, estimating the MI between one continuous and one discrete random variable is in general non-trivial and multiple approaches exist for this estimation, such as nearest-neighbor^24^ and binning methods. To enable our MI calculations (such as *I*(*r* ; *D*)), we used a procedure that bins data points of *r* and avoids biased MI estimates in our finite but large sample size. In computing MI, we treated the binned feature vector 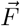 (where 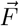 has the possibility of being one dimensional when representing *r* or *n*) and the degree of relatedness *D* as two discrete random variables with realizations *f* and *d* ∈ [1, 7], respectively. If we know the probability mass functions (pmfs) of the discrete random variables *X* and *Y* with realizations *x* and *y*, we can calculate MI using its definition as

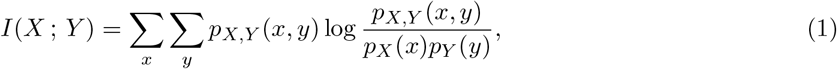

where *p*_*X,Y*_ is the joint pmf of *X* and *Y* and *p*_*X*_, *p*_*Y*_ are the marginal pmfs of *X* and *Y* respectively.

Binning a continuous variable in order to use Equation (1) introduces the difficulty of picking the right bin size. It has been shown^24^ that MI is sensitive to bin size and that its stability with respect to this variable is dependent on the sample distribution. Our distributions and sample sizes of *r* yielded a large range of bin sizes that have stable MI estimates (see the flat regions of each curve in Figure S7). Because the fraction *G*_*N*_ is normalized by MI, its correct calculation relies on the unbiasedness of the various MI quantities that form it. At bin sizes smaller than 150 pairs per bin (ppb), both the means and standard deviations (Figure S8) of our MI quantities increased rapidly. Given this, in our calculations of MI, we binned *r* at 150 ppb, where our binning converts a continuous value of *r* to its nearest bin-value in 150 evenly spaced numbers from 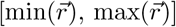. Here and below 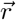 represents all sampled training and testing data points *r*.

### Estimating mutual information

Calculating Equation (1) without access to the entire spaces *𝒳* and *𝒴*—i.e., estimating MI from sampled data—is contingent on the estimation of marginal and joint probabilities *p*_*X*_, *p*_*Y*_, and *p*_*X,Y*_. We used a simple counting approach to calculate each probability, assigning 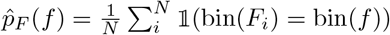. Here 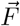 is the vector of realized data points representing all *N* sampled values for the desired feature *r, n*, or (*r, n*); bin(*x*) denotes the function that takes a continuous realization to its binned value; and 𝟙(*X* = *Y*) is the indicator function. By binning *r* to 150 ppb as noted in the previous subsection, we were able to use this discrete maxium likelihood estimator (MLE) approach for calculating every desired pmf and obtain stable results in MI.

We performed calculations of MI on the exact IBD segment data restricted to three distribution shapes: A uniform distribution, a “slow-exponential” distribution, and an exponential distribution (see Figure S1). We accounted for different distributions of *D* in the calculations of 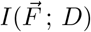 by decomposing the joint pmf relating 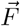 and *D* as *p*_*F*_ (*f, d*) = *p*_*F*_ (*f* |*d*)*p*_*D*_(*d*), and also decomposing the marginal pmf on 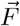 (with the law of total probability *p*_*F*_ (*f*) = ∑_*d′*_ *p*_*F*_ (*f* |*d′*)*p*_*D*_(*d′*)). Equation (1) is then expressed as

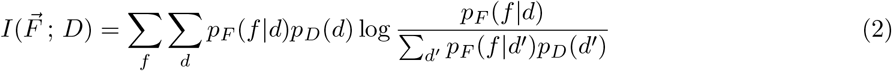

by canceling the *p*_*D*_(*d*) terms in the numerator and denominator. *p*_*F*_ (*f* |*d*) is the pmf of realizations of feature 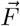 in a given degree, and *p*_*D*_(*d*) is the distribution shape (from Figure S1). This approach removes noise associated with calculating the pmfs 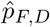 for different distribution shapes, which stems in part from random factors in finite sample sizes (including smaller numbers of pairs in the non-uniform distributions). In particular, because the probabilities of *f* conditioned on any given degree of relatedness *d* are identical across each distribution shape, we estimated 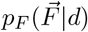 once only from the uniform distribution data. Note too that the differences in MI due to the *D* distribution are entirely accounted for in the probabilities *p*_*D*_(*d*), and these are exactly calculable given the equation for each distribution.

### Probability density estimation of features

In the context of the Bayes classifier, we estimated the probability of a feature realization *f* conditioned on the training data 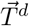 in degree *d* according to the degree-wise count as

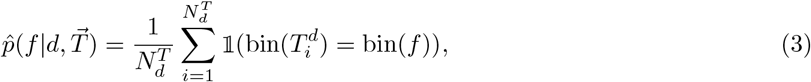

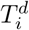 being a particular realization of the training data 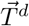 with total count 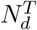. However, we only had access to the frequencies of realizations *f* that occur at least once in the training data, so Equation (3) is only calculable for these values. The total training data 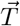 and testing data 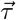 are of dimension equal to their respective number of data points *N*^*T*^ or *N*^*τ*^. To generate posteriors *p*(*D*|*τ*_*i*_) for realizations in 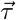 at values where there are no training data points in bin(*τ*_*i*_), we linearly interpolated the values given by Equation (3) within the convex hull (see Figure S9) specified by the bounds of the training data. (Strictly speaking, these posteriors are then incorrect pmfs with mass greater than 1—however, in practice this is only relevant for a vanishingly small number of points.) We used the `scipy` packages `interp1d` and `griddata` for the linear interpolations in one-dimension (when 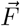 is either *r* or *n*) and two-dimensions (when 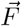 is (*r, n*)), respectively.

In the case that 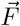 is (*r, n*), the two-dimensional linear interpolation of 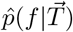 values are only well-defined inside of the two-dimensional convex hulls of the training data. Therefore, we could not assign posteriors to realizations of the testing data that lay outside the bounds of the training data. For these data points (labeled in Figure S9 as “Unscored under (*r, n*)”), we assigned probability values according to the one-dimensional interpolation of 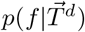 values with 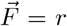. The one-dimensional interpolations for 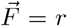 (or 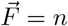) only remained undefined when they occurred outside the interval of training values 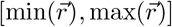 (or 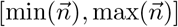), in which case they remained unclassified in our analysis. In the inferred segment data, there was only a maximum of one point per degree that remained unclassified.

### Bayes classification

Our classifiers use the posterior probabilities 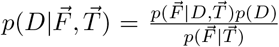 for the single and multivariate features 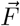 to infer *D* in the testing data. The priors *p*(*D*) are the known shape of the degree distribution (Figure S1), and we generated the probability of our data 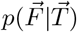 as the sum across degrees according to the law of total probability 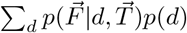. We calculated likelihoods 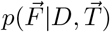 according to the estimator in Equation (3). To classify a testing pair *τ*_*i*_ to a certain degree, we calculated log *p*(*D*|*τ*_*i*_) for each degree and classified the pair as the maximum a posteriori degree:

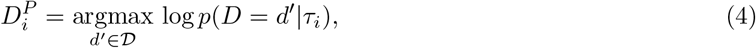

where 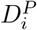 is the predicted degree, and *𝒟* is the set of possible degrees *{*1, …, 7*}*. The recall of a particular classifier for degree *d* is 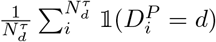.

The classifier takes input IBIS segments and calculates the IBD proportion and segment numbers for all pairs of individuals with at least one detected IBD segment. It classifies any pair with *r <* 2^−15*/*2^ (a common lower bound for seventh degree classification^11,14^) as unrelated, and, for all other pairs, predicts their degree using Equation 4.

### Simulated data

For the exact IBD segment data, we used Ped-sim^22^ to simulate 231,000 relative pairs of 13 relationship types from seven degrees of relatedness (Table 1) (replicated 80 times for the MI analysis and once for the classification-based analysis) and leveraged the IBD segments this tool prints. Thus these segments are free of error and we refer to them throughout as *exact*. We used both sex-specific genetic maps^25^ and crossover interference modeling^26^ for these simulations.

For each degree, we simulated an equal number of pairs from each of two relationship types. The one exception is first degree relatives where we only considered full sibling pairs since parent-child pairs always have *r* = 0.5 and are trivial to identify. We doubled the number of full sibling pairs (to the total number assigned from the distribution shape) so that the first degree relatives included the full number of pairs. We calculated the IBD proportion by adding the lengths of all outputted IBD segments and dividing by the total length of the sex averaged genetic map—halving the length of IBD1 segments (see the equation for *r* in Results). We calculated the segment number by counting the number of outputted IBD segments from either Ped-sim (exact) or IBIS (inferred, as described next).

To simulate relatives with genetic data, we used autosomal genotypes from participants in the UK Biobank^2^ as founders in Ped-sim runs. We used the phased data distributed by the UK Biobank^2^ and, before simulating, filtered the samples to include the white British ancestry subjects. To filter out close relatives, we first performed SNP quality control filtering on the UK Biobank unphased genotypes (filtering SNPs with minor allele frequency less than 2%, missing data rate greater than 1%, and retaining only SNPs used for phasing in the original analysis^2^). Next we ran IBIS v1.20.8 on the filtered genotypes with the `-maxDist 0.12` option and with IBD2 segment detection enabled. This provided kinship coefficients that we then input to PRIMUS^3^, running it with `--no_PR` (which corresponds to not reconstructing pedigrees: executing only IMUS^27^) and `--rel_threshold 0.022` to filter out relatives with a kinship coefficient greater than 0.022 (i.e., retaining only pairs no more closely related than fifth degree^11^). We ran Ped-sim as described above (using sex-specific genetic maps and crossover interference modeling) and otherwise used default options (including genotyping error and missing data rates of 10^−3^). Finally, we used IBIS v1.20.7 (enabling IBD2 detection with `-2`) to detect IBD segments between these simulated relatives.

### Running ERSA

To get relatedness estimates from ERSA^15^, we first ran GERMLINE^23^ v1.5.1 with `-err_het=1` and `-err_hom=2` (the options recommended by the ERSA authors) on the simulated Ped-sim haplotypes. That is, we provided ERSA perfectly phased data output by the simulator. We then ran ERSA with default options on the resulting GERMLINE segments.

### Runtimes

We ran both ERSA and our Bayes classifier on a machine with an AMD EPYC 7702 2.0 GHz processor and 1 TB of RAM. We supplied 16GB to ERSA and 8GB to our Bayes classifier. Both methods are single threaded.

## Discussion

In this paper, we sought to examine how much incorporating the number of IBD segments together with the coefficient of relatedness of a relative pair improves degree of relatedness inference. We thus provided both a theoretical MI analysis using simulated exact IBD segments and a machine learning-based classification analysis using exact and inferred segments. The results using exact segments show that including IBD segment numbers can non-trivially enhance related inference quality, especially for distant relatives. However, the results using inferred segments reveal that IBD detection errors—including false negatives for segments shorter than 7 cM—meaningfully limit this improvement. Indeed, the performance of our machine learning classifier is almost indistinguishable from IBIS (Figure 3), which does not use segment numbers. With the potential development of more accurate IBD detection tools in the future—including for whole genome sequencing data—use of IBD segment numbers in relatedness inference models may be worth considering.

**Figure 3:**
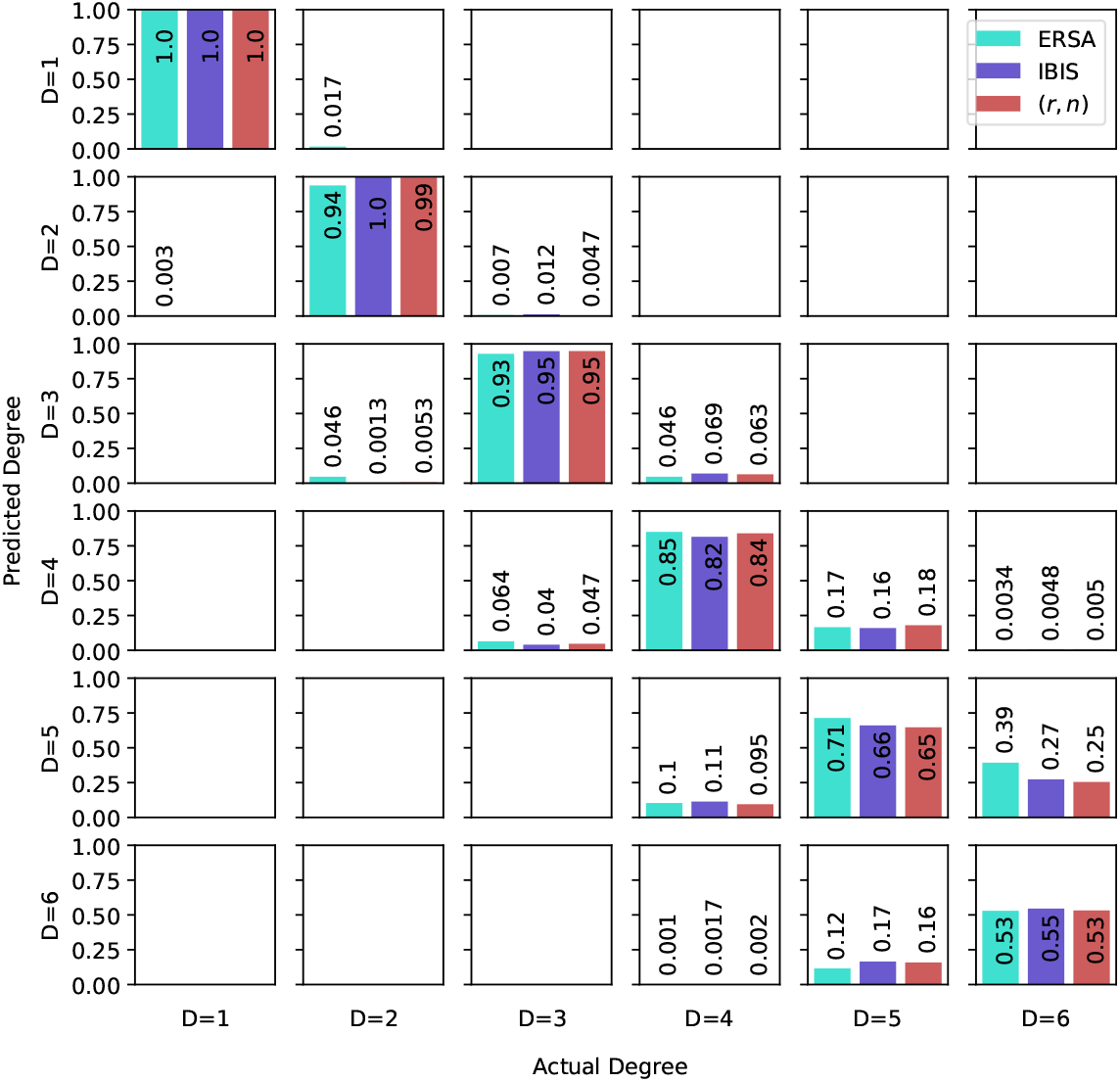
Confusion matrix of recalls of ERSA, IBIS, and our Bayes classifier trained on (*r, n*).

We introduced a machine learning-based classifier and demonstrated that it has comparable accuracy to two state-of-the-art methods and is computationally efficient. Because we fit the classifier to population-specific training data (instead of using fixed kinship thresholds for each degree^12,14^), it implicitly accounts for background IBD sharing and erroneous IBD signals. This approach differs from model-based methods such as ERSA in that it makes no assumptions about the distributions of IBD segment lengths or numbers with respect to relatedness degrees. Those assumptions can be violated in populations with small effective size or a historical founder effect^15^. Our trials of this machine learning method suggest that even without data for large numbers of (labeled) real relatives, simulating relatives enables this data-driven approach to relatedness inference. Additionally, both the machine learning classifier and the MI analyses can be easily extended to include other IBD features such as the minimum or maximum IBD segment length between a pair.

An important factor in attempting to utilize IBD segment numbers is their accurate detection. Switch errors profoundly influence segment number estimates when using phase-based IBD detectors^12,13,16,28^. Our use of IBIS segments in our classifier was motivated by IBIS’s ability to call IBD segments in unphased data— one of only a few methods to do so^13^—which is key to avoiding biased segment number estimates. ERSA takes inferred IBD segments from the phase-based IBD detector GERMLINE. To exclude the possibility of phasing errors impacting ERSA’s performance, the phased data we provided GERMLINE was that generated by the simulator, thus being perfect up to the limit of the haplotypes input to Ped-sim. In particular, these haplotypes do not contain switch errors in IBD segments between the simulated relatives. It is possible that ERSA’s superior performance in classifying fifth degree relatives is enhanced by its segment detection in these data.

In general, our analyses are consistent with prior work showing that relatedness inference can achieve high recall for up to third degree relatives. However, two recent papers have focused on distinguishing relationship types of the same degree, especially three types of second degree relatives^29,30^. In this setting, IBD segment numbers can provide useful information, such as for distinguishing avuncular from grandparent-grandchild pairs^1^. Still, for degree of relatedness inference, even when using exact IBD segments, the classification recalls for distant relatives—i.e., those beyond fourth degree—are limited (Figure 2(a)). This suggests that pairwise IBD information might not be sufficient to reliably infer distant relatives, regardless of segment quality. Approaches that leverage multi-way IBD signals to infer more distant relatives can achieve considerably higher accuracy than those of pairwise methods^31,32^. Even so, these multi-way methods are built on pairwise classifiers, so understanding and improving pairwise relatedness classification remains an important fundamental problem for relatedness inference.

## Acknowledgements

We thank Debbie Kennett for conversations about genetic genealogy. Funding for this work was provided by NIH grant R35 GM133805. Computing was performed on a cluster administered by the Biotechnology Resource Center at Cornell University. This research has been conducted using the UK Biobank Resource under Application Number 19947.

## Declaration of Interests

A.L.W. is a paid consultant for 23andMe and the owner of HAPI-DNA LLC. The other authors declare no competing interests.

## Notes

### Summary of Updates

Incorrect author order in original version. Jesse Smith, Ying Qiao, Amy L. Williams is correct.

