## Supplementary Figures for "Evaluating the utility of identity-by-descent segment numbers for relatedness inference via information theory and classification"

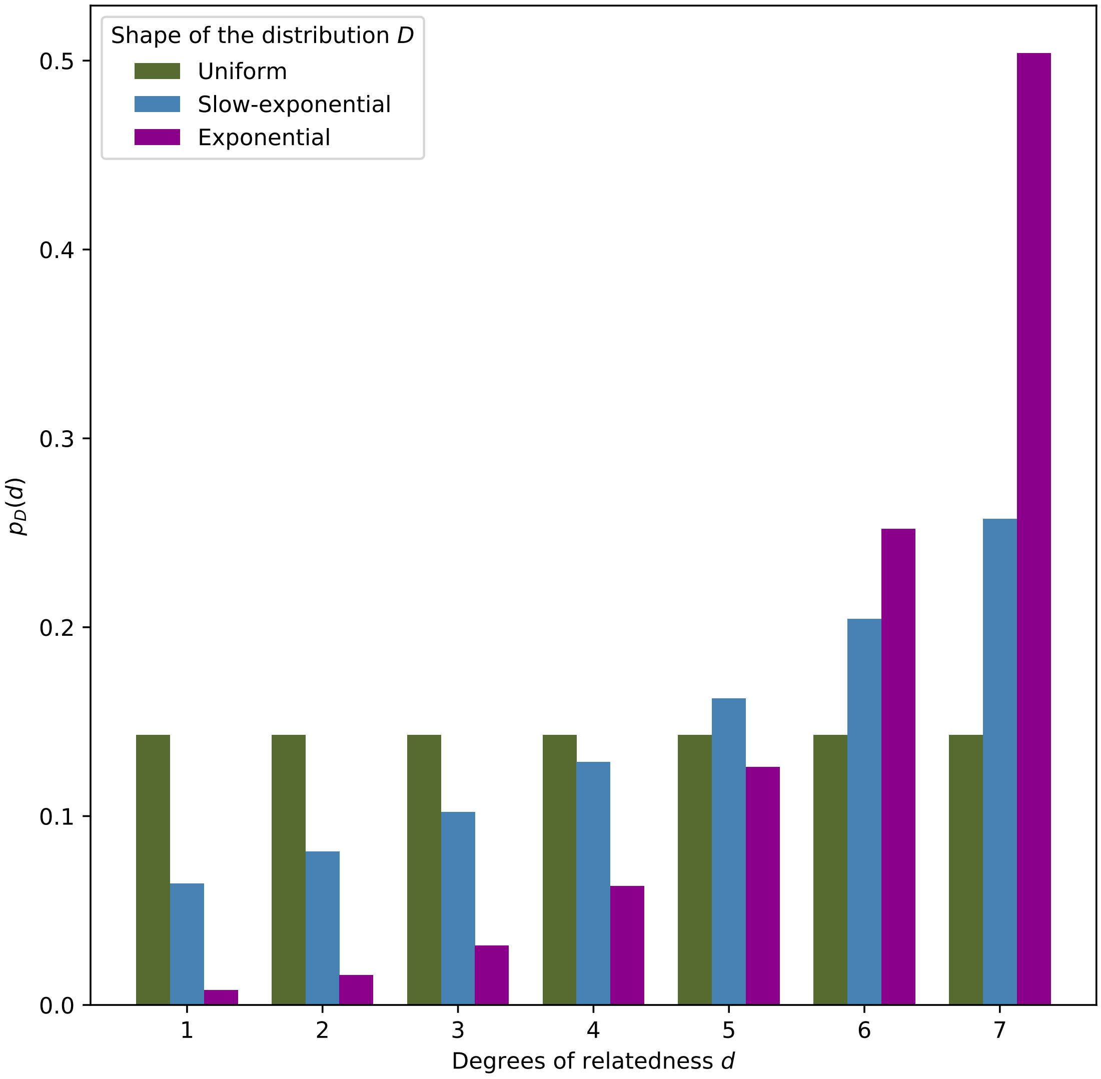


**Figure S1:** Probability mass functions of different distribution shapes for $D$ as a function of degree of relatedness $d$, where uniform=1/7, slow-exponential=(1000/15541) $\times$ $2^{(d-1)/3}$, and exponential=(160/20320) $\times$ $2^{d-1}$. Total pair counts for the testing data are 21,000 for uniform, 15,541 for slow-exponential, and 20,320 for the exponential.

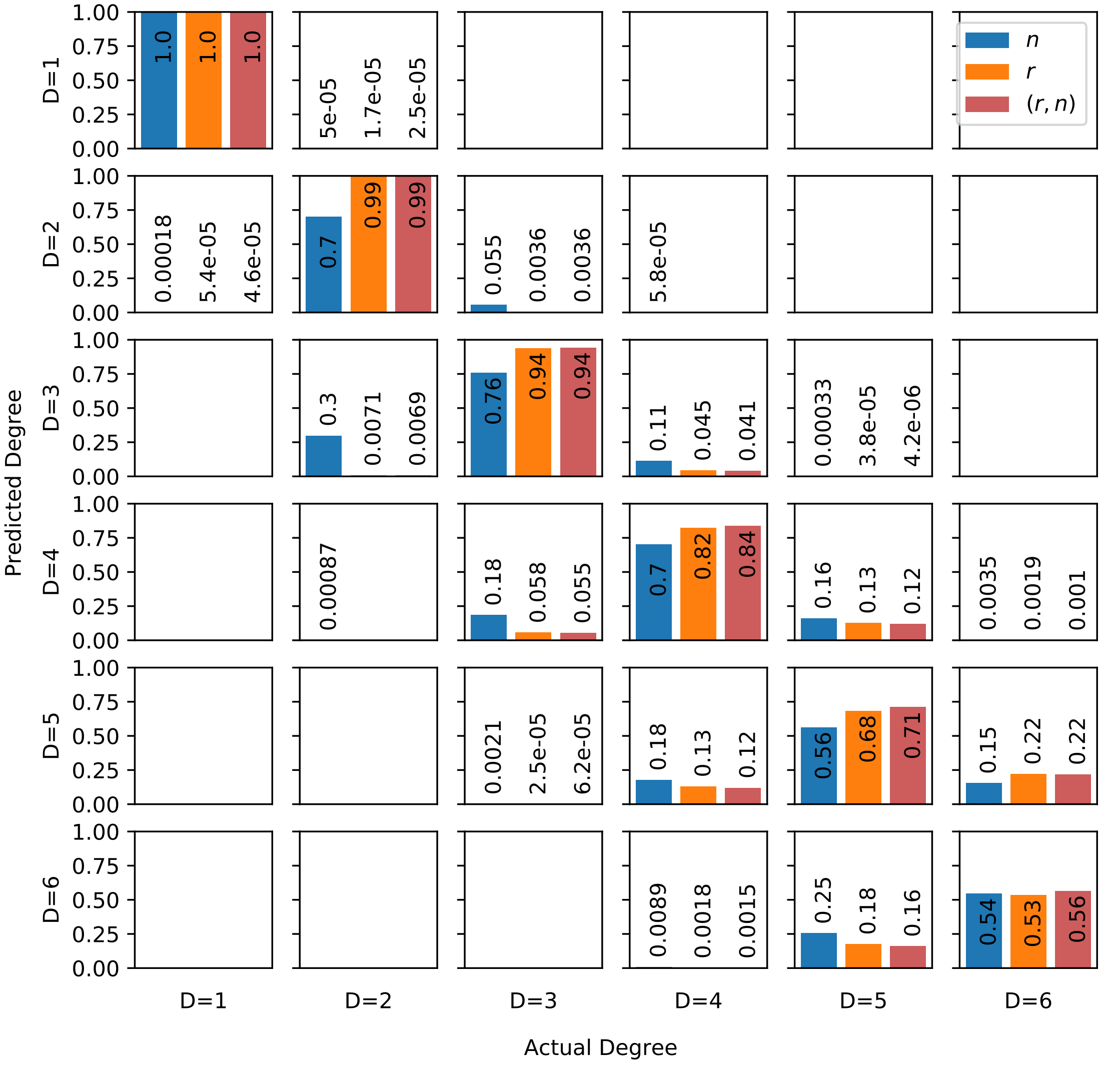


**Figure S2:** Confusion matrix (with respect to degree of relatedness) of Bayes classifiers trained on exact segments with features $n$, $r$ and $(r, n)$ from the uniform distribution. Most misclassifications occur in diagonal-adjacent cells (off-by-one-degree misclassifications).

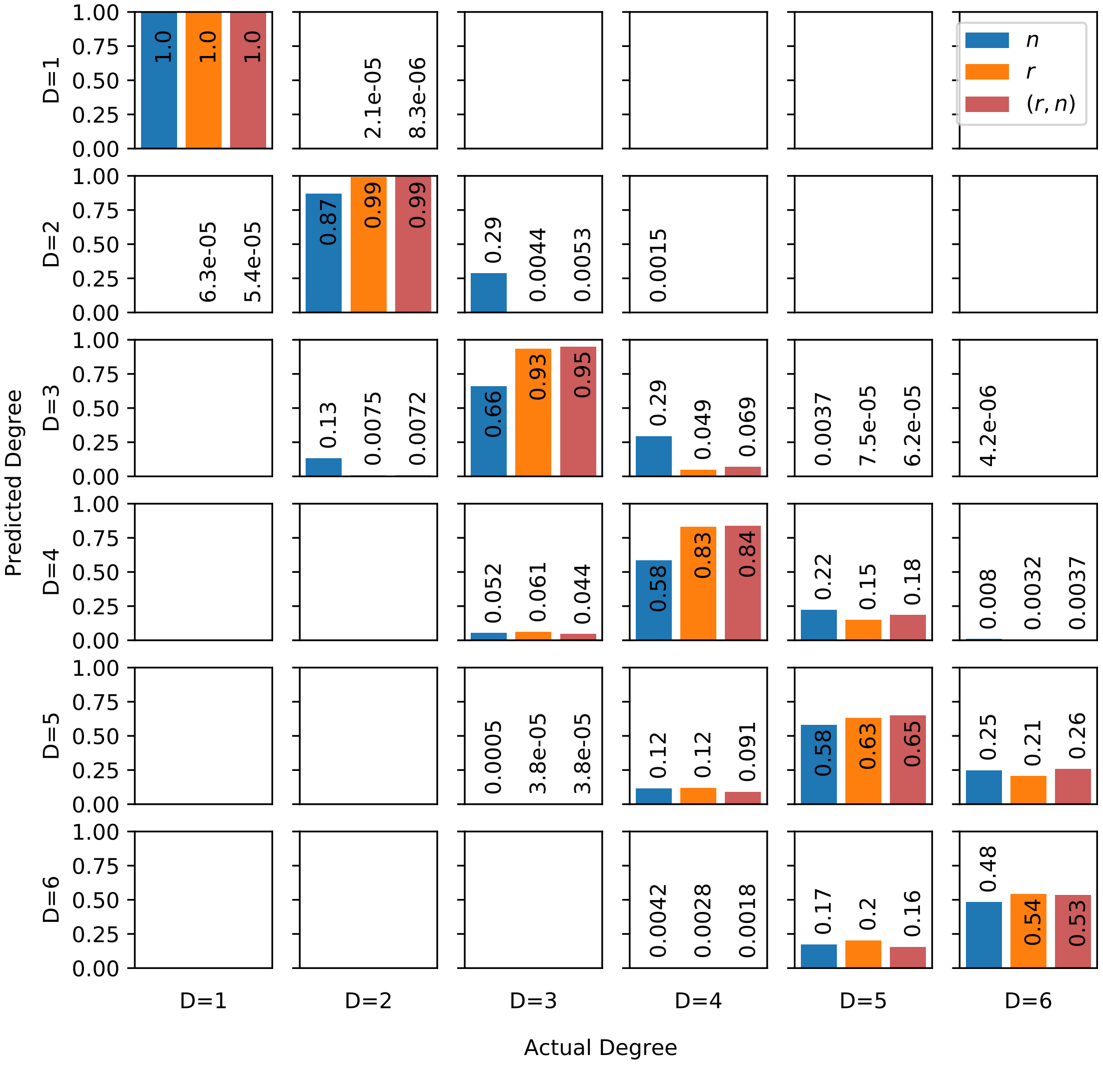


**Figure S3:** Confusion matrix (with respect to degree of relatedness) of Bayes classifiers trained on inferred segments with features $n$, $r$ and $(r, n)$ from the uniform distribution. Most misclassifications occur in diagonal-adjacent cells (off-by-one-degree misclassifications).

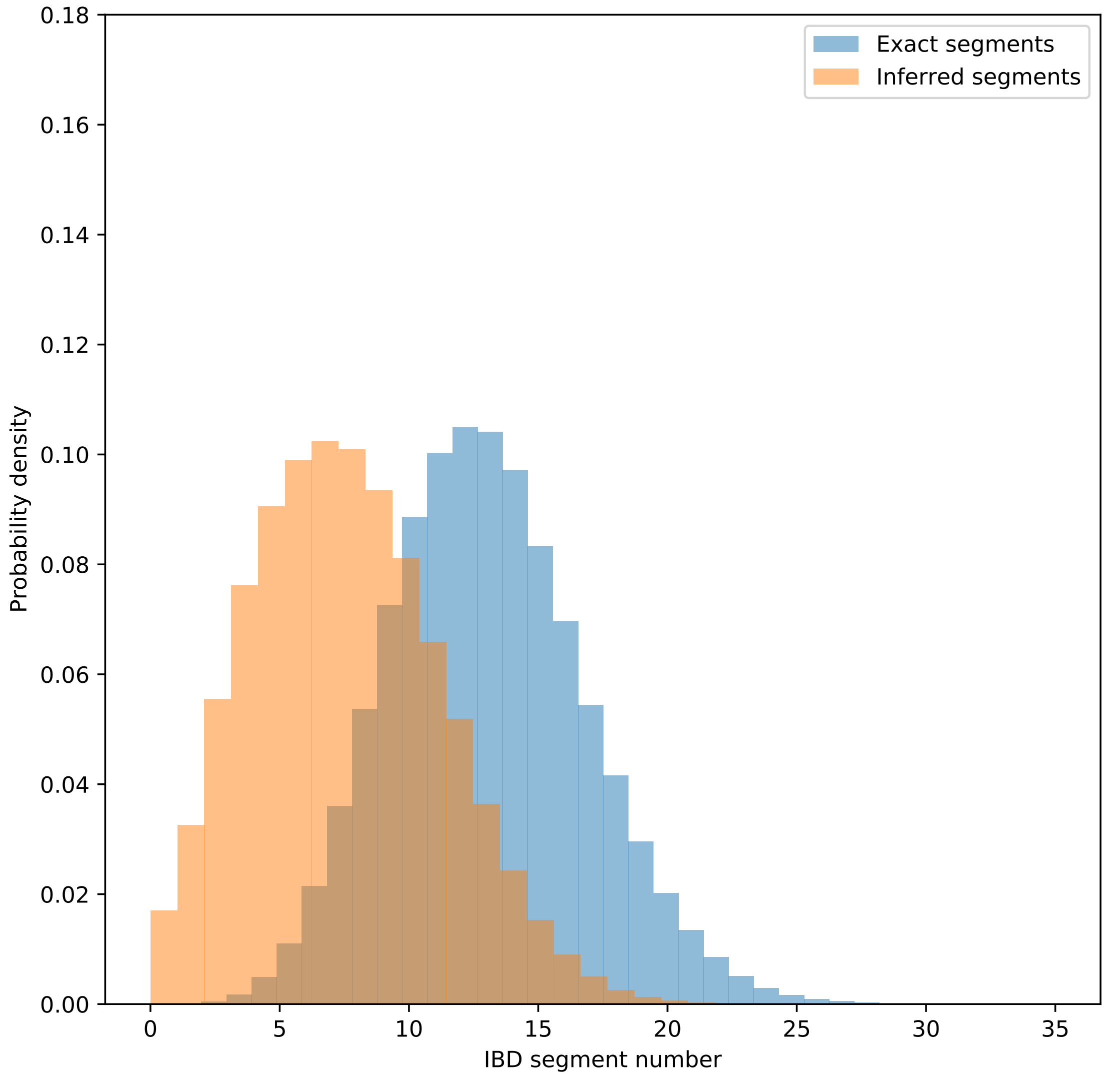


**Figure S4:** Distributions of exact and inferred segment numbers in fifth degree pairs.

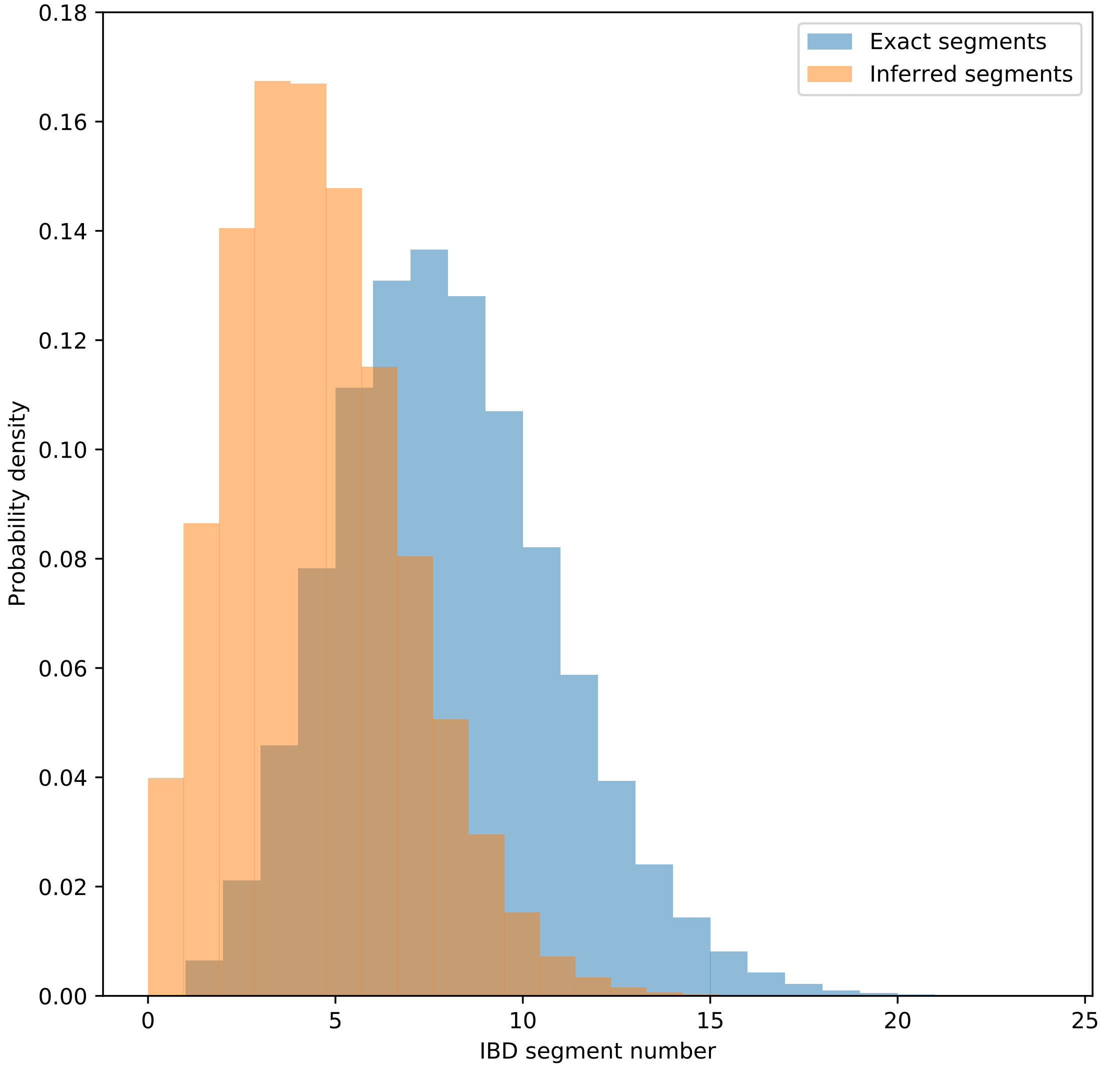


**Figure S5:** Distributions of exact and inferred segment numbers in sixth degree pairs.

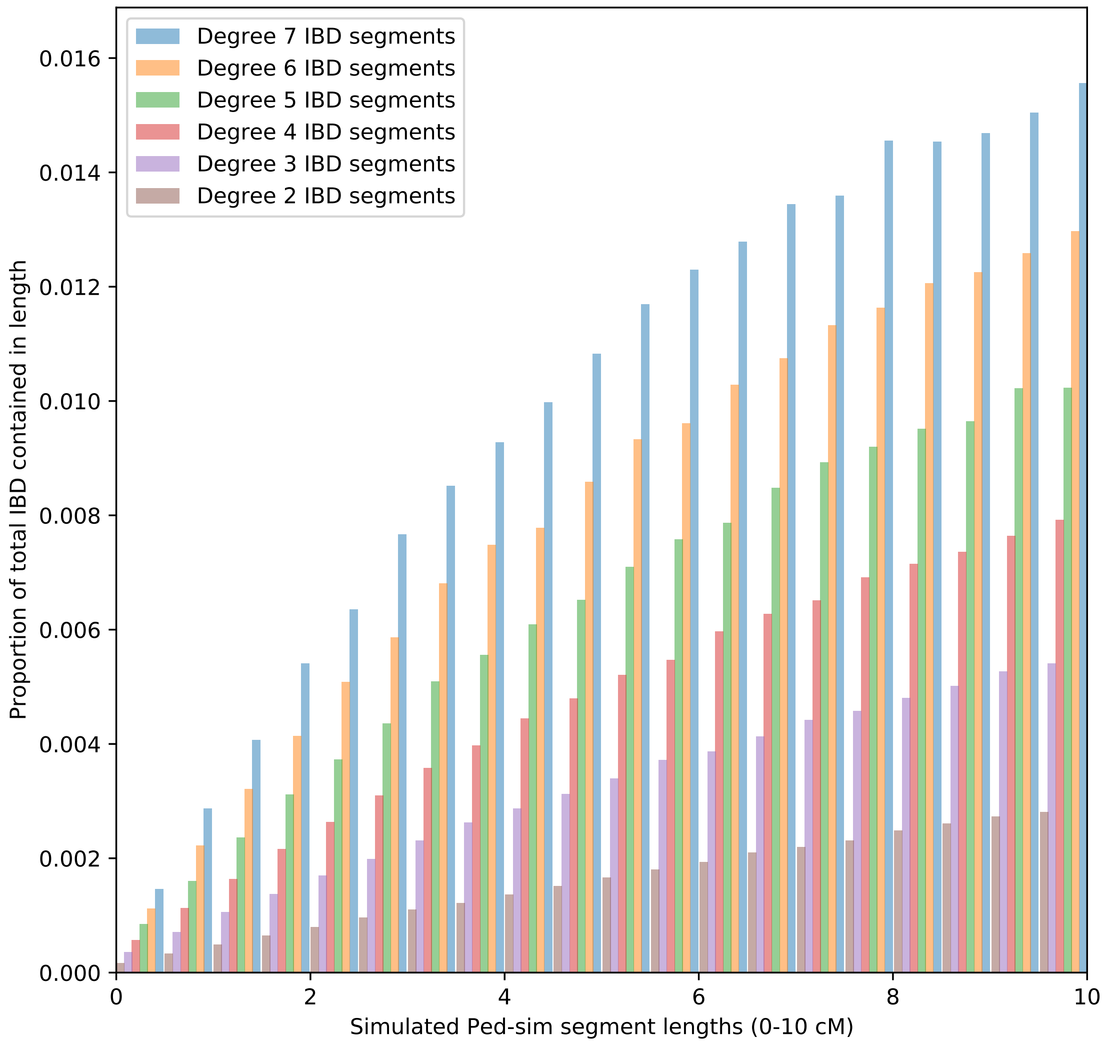


**Figure S6:** Average proportions of pairwise total IBD length contained in exact segments of lengths 0-10 cM for relatives of the indicated degrees. Proportions calculated over 33,000 pairs from each degree.

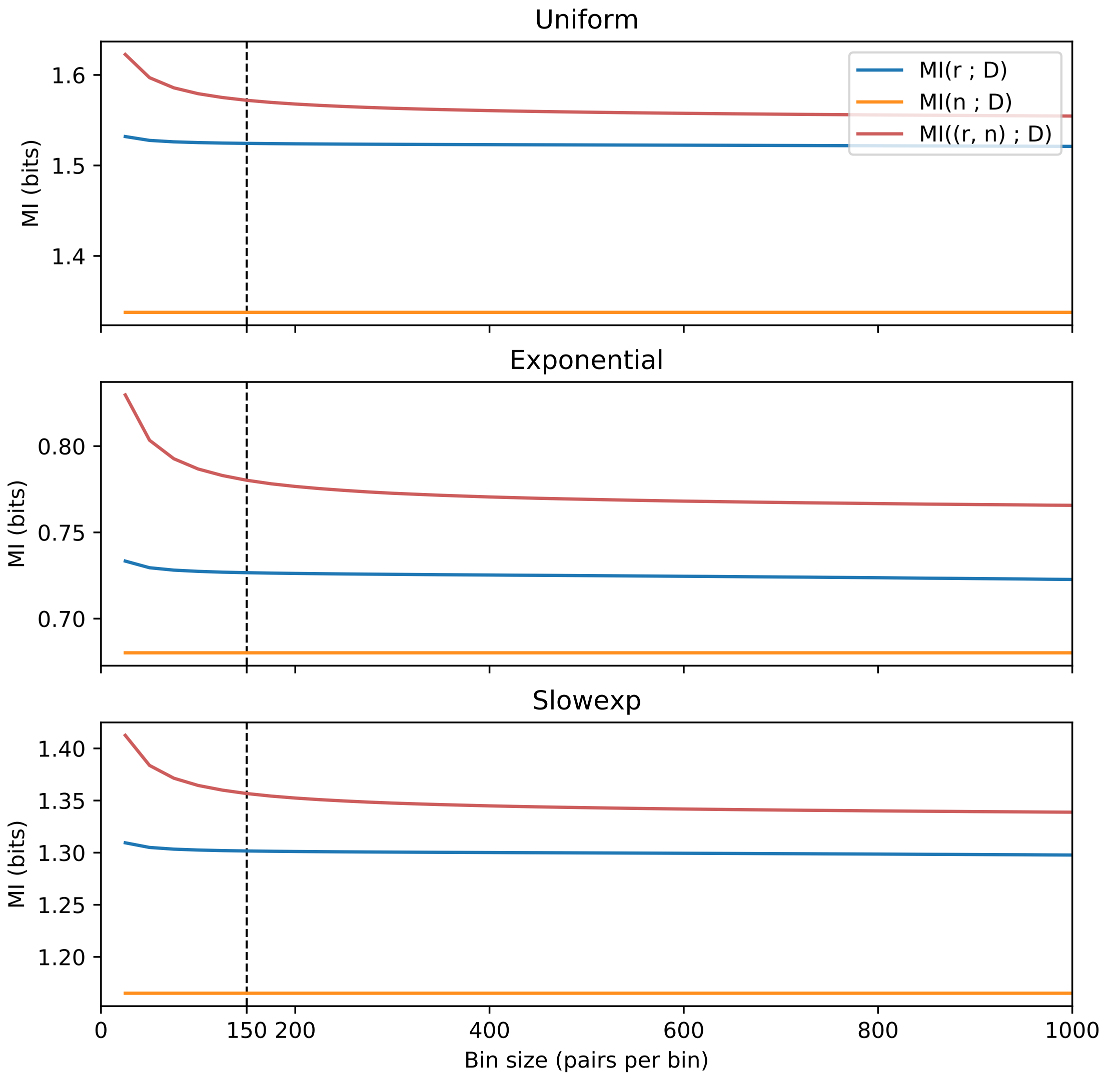


**Figure S7:** MI of different feature sets as a function of bin size (pairs per bin), averaged over 80 independent simulations of exact segments from each of the distribution shapes. Slowexp corresponds to the slow-exponential distribution.

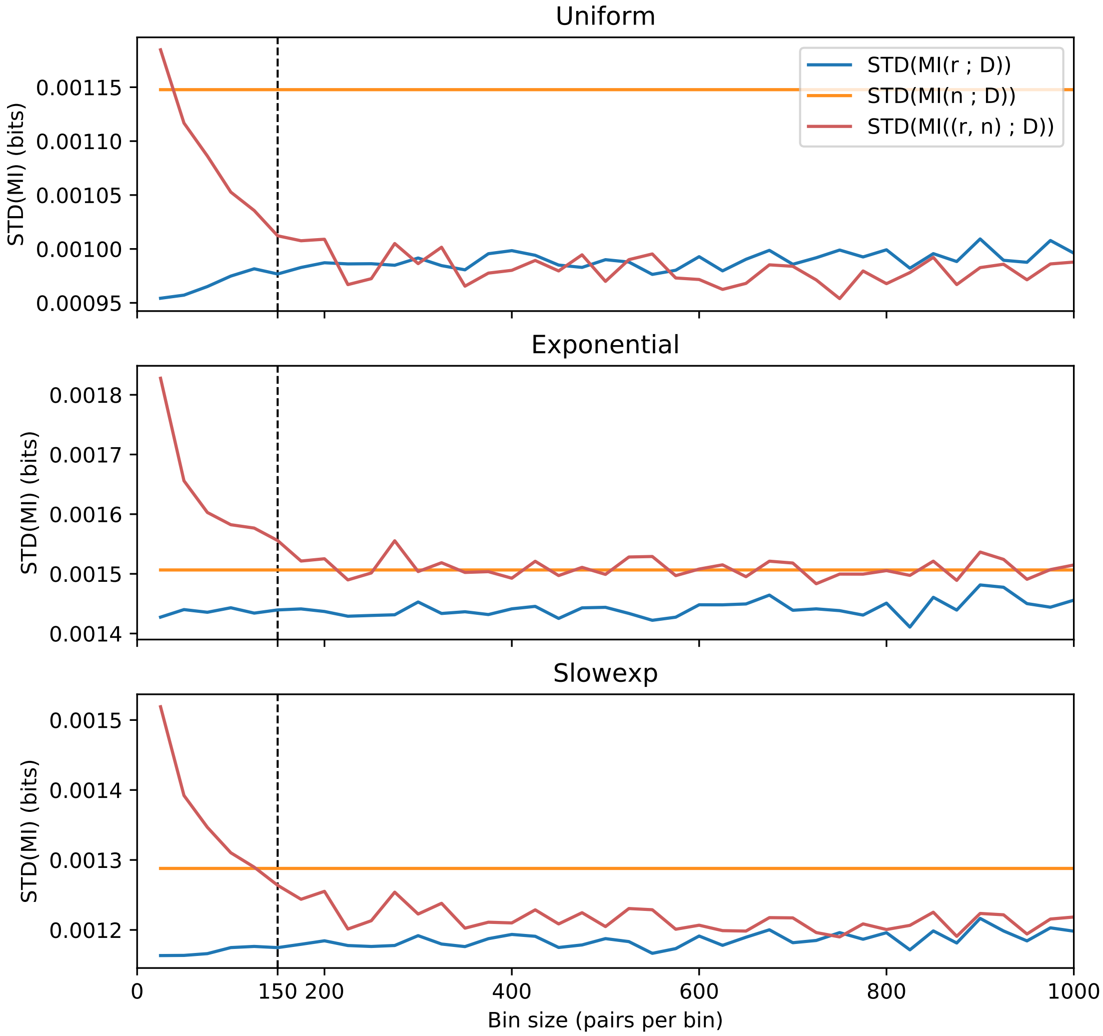


**Figure S8:** Standard deviations of MI of different feature sets as a function of bin size (pairs per bin), averaged over 80 independent simulations of exact segments from each of the distribution shapes. Slowexp corresponds to the slow-exponential distribution.

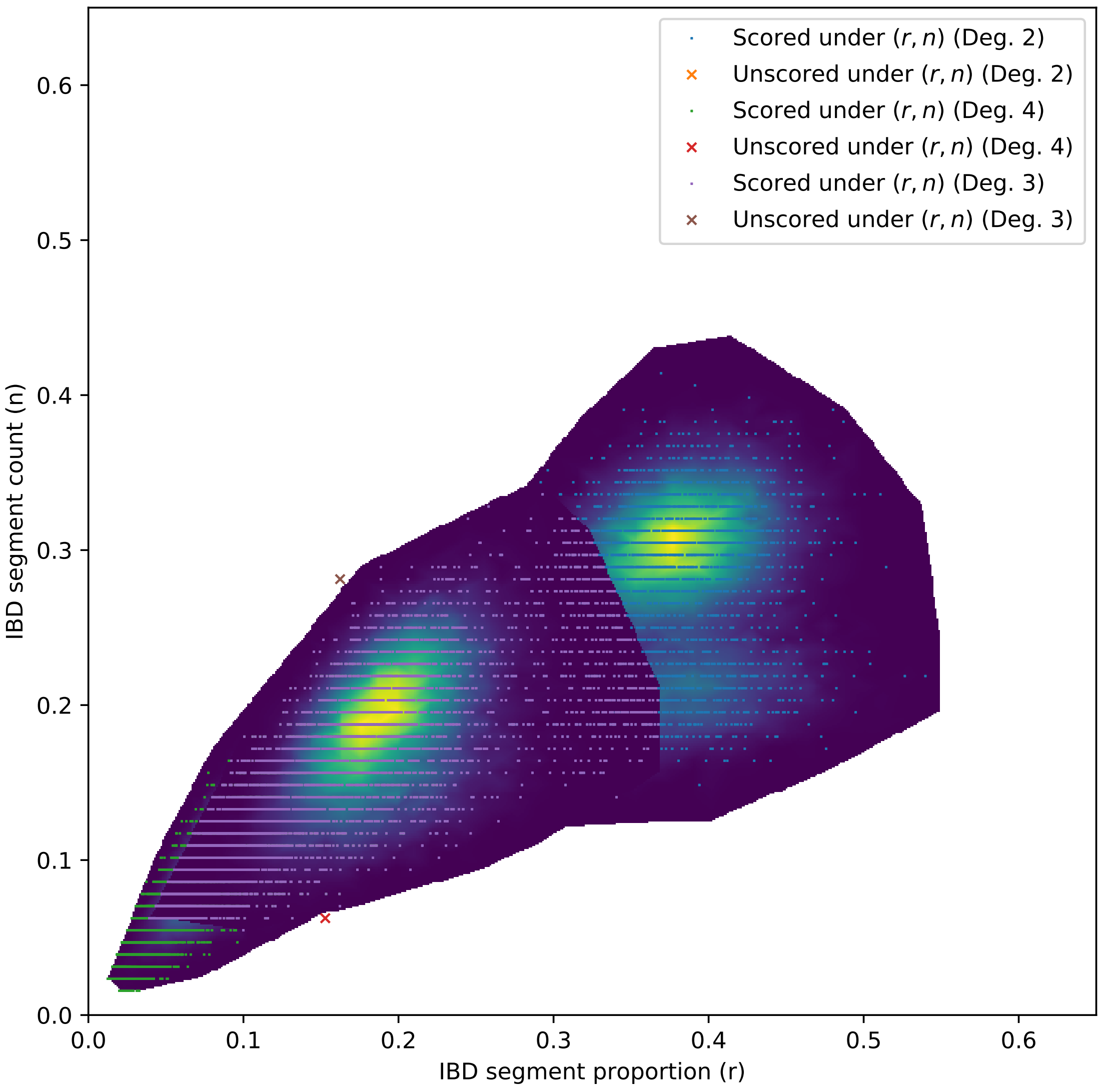


**Figure S9:** Heat maps depicting posteriors $\hat{p}(D|f,\vec{T})$ for inferred IBD segments for several values of $D$. Generated using the `griddata` two-dimensional interpolation on $\hat{p}(f|D)$ calculated from training data. Overlaid are corresponding testing data points colored by their classification. Here, the IBD segment number $n$ has been normalized to unity. Probabilities and points from higher degrees are plotted on top of those from lower degrees.
